# Overestimated Polygenic Prediction due to Overlapping Subjects in Genetic Datasets

**DOI:** 10.1101/2022.01.19.476997

**Authors:** David Keetae Park, Mingshen Chen, Seungsoo Kim, Yoonjung Yoonie Joo, Rebekah K. Loving, Hyoung Seop Kim, Jiook Cha, Shinjae Yoo, Jong Hun Kim

## Abstract

Recently, polygenic risk score (PRS) has gained significant attention in studies involving complex genetic diseases and traits. PRS is often derived from summary statistics, from which the independence between discovery and replication sets cannot be monitored. Prior studies, in which the independence is strictly observed, report a relatively low gain from PRS in predictive models of binary traits. We hypothesize that the independence assumption may be compromised when using the summary statistics, and suspect an overestimation bias in the predictive accuracy. To demonstrate the overestimation bias in the replication dataset, prediction performances of PRS models are compared when overlapping subjects are either present or removed. We consider the task of Alzheimer’s disease (AD) prediction across genetics datasets, including the International Genomics of Alzheimer’s Project (IGAP), AD Sequencing Project (ADSP), and Accelerating Medicine Partnership - Alzheimer’s Disease (AMP-AD). PRS is computed from either sequencing studies for ADSP and AMP-AD (denoted as rPRS) or the summary statistics for IGAP (sPRS). Two variables with the high heritability in UK Biobank, hypertension, and height, are used to derive an exemplary scale effect of PRS. Based on the scale effect, the expected performance of sPRS is computed for AD prediction. Using ADSP as a discovery set for rPRS on AMP-AD, ΔAUC and ΔR^2^ (performance gains in AUC and R^2^ by PRS) record 0.069 and 0.11, respectively. Both drop to 0.0017 and 0.0041 once overlapping subjects are removed from AMP-AD. sPRS is derived from IGAP, which records ΔAUC and ΔR^2^ of 0.051±0.013 and 0.063±0.015 for ADSP and 0.060 and 0.086 for AMP-AD, respectively. On UK Biobank, rPRS performances for hypertension assuming a similar size of discovery and replication sets are 0.0036±0.0027 (ΔAUC) and 0.0032±0.0028 (ΔR^2^). For height, ΔR^2^ is 0.029±0.0037. Considering the high heritability of hypertension and height of UK Biobank, we conclude that sPRS results from AD databases are inflated. The higher performances relative to the size of the discovery set were observed in PRS studies of several diseases. PRS performances for binary traits, such as AD and hypertension, turned out unexpectedly low. This may, along with the difference in linkage disequilibrium, explain the high variability of PRS performances in cross-nation or cross-ethnicity applications, i.e., when there are no overlapping subjects. Hence, for sPRS, potential duplications should be carefully considered within the same ethnic group.

## INTRODUCTION

Recently, genetic studies involving a polygenic risk score (PRS) have dramatically grown in number, and sophisticated tools and methodologies are being developed for its use^1–3^. Along with heritability, PRS has become an important metric for explaining complex genetic diseases (e.g. Alzheimer’s disease, AD) and traits^4–8^. A typical PRS study involves both the discovery and replication phases^9,10^. In the discovery phase, two different methods are used to develop PRS. PRS is essentially derived from the raw genetic data, denoted as rPRS. Instead, the summary statistics from large-scale genetics studies or GWAS catalogs may be also be used, which we abbreviate as sPRS. Polygenic prediction performance is then evaluated by the marginal contribution of the PRS term in a regression model on target clinical application^10,11^.

An underlying assumption of PRS models is that the subjects from the discovery set do not overlap with those of the replication set^10^. However, our preliminary analyses (Fig. 1a) demonstrate a significant number of identical subjects across multiple genetic datasets. The overlapping subjects may be identified and removed for rPRS using the raw genetic data, a challenge remains for sPRS in which raw data is inaccessible. Therefore, we posit that a strict level of independence across datasets is hard to achieve with sPRS. This may pose serious issues to related fields, since the subject-level dependence across datasets may not only inflate the polygenic prediction performance, but also prevent generalizable applications of the developed model.

**Fig. 1.**
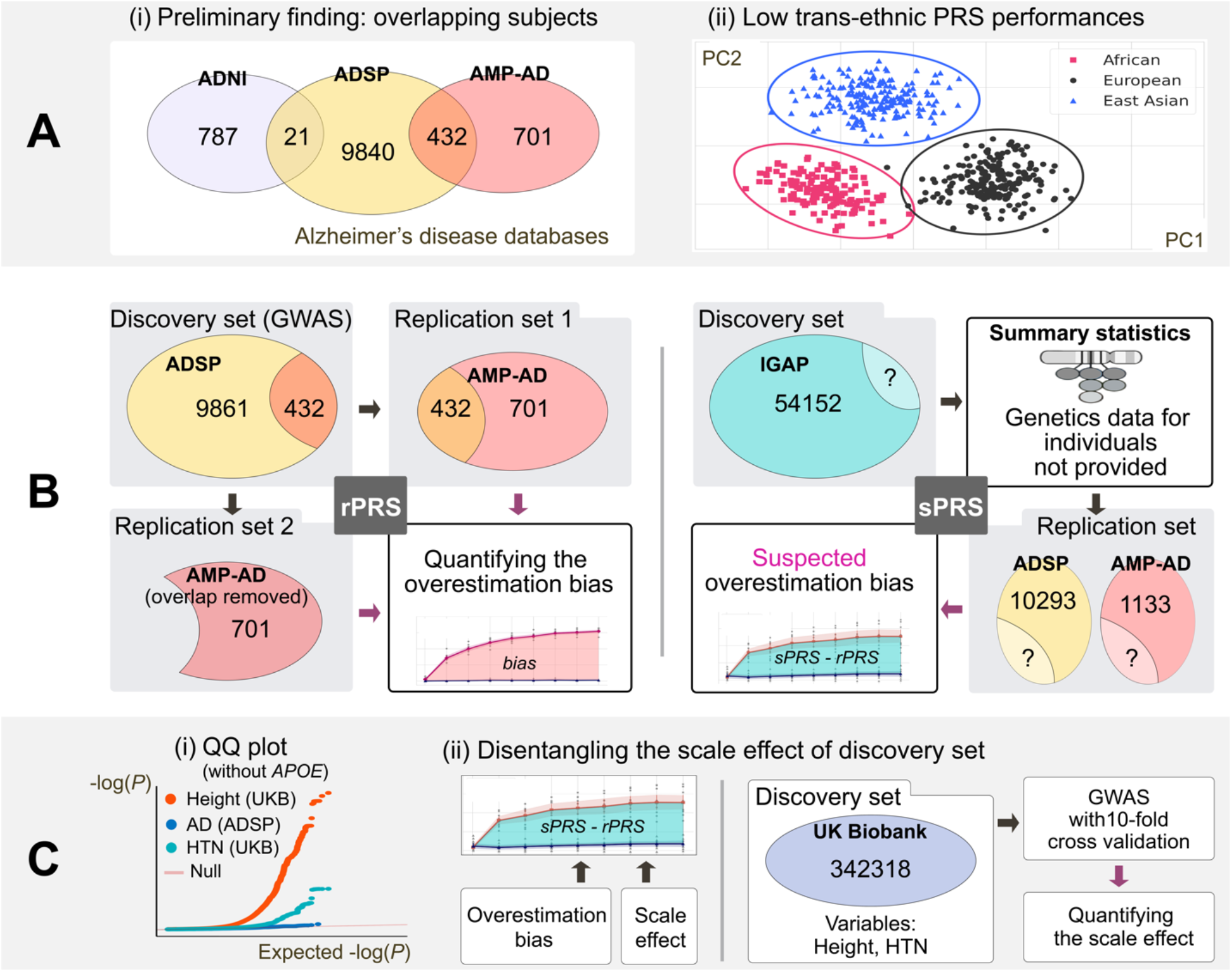
Overview of the study. (A) (i) Overlapping subjects are observed between AD genetic initiatives. (ii) Without a direct comparison, there is no overlapping subject across ethnicities. Until now, trans-ethnic applications of PRS have been limited. We suspect that overlapping subjects within an ethnicity are one of the key factors to explain overestimated performances, which motivates this study. We divide PRS into two cases, where rPRS represents when the genetic information is provided and used as the discovery set and sPRS stands for the case when GWAS is pre-conducted and only summary statistics are provided. (B) For rPRS, overlapping subjects (n=432) between ADSP and AMP-AD are identified, which breaks the independence assumption and causes the overestimation bias. For sPRS, the overlapping ratio cannot be examined given the summary statistics. However, the suspected inflation in the AD prediction performance (denoted by sPRS - rPRS) motivates further analysis into the scale effect of the datasets because IGAP has a larger number of samples. (C) (i) Two new variables, hypertension and height, from the UK Biobank database are introduced to compute the upper bounds of the scale effect. Hypertension and height have a higher heritability than AD. Thus, they act as the upper bounds for AD over PRS performances (shown in the QQ plot). (ii) In AD, the gap between sPRS and rPRS (area shaded in green) is attributable to either the overestimation bias or the scale effect of the sample size of the discovery set. Because UK Biobank consists of a larger number of samples (n=342318), the scale effect can be measured via computing the performance gains per sample unit.

Among prior studies, we identify multiple signs of potential inflation in PRS performance attributable to overlapping subjects. First, if the independence between discovery and replication sets is clearly stated in the paper, the PRS performance of binary traits not statistically significant, while if not, results were highly variable or inflated^12–16^. For instance, for models in which independence is explicitly controlled during development, even with a large-scale national biobank, PRS contributed less than 2% in the model accuracy^17^. As a similar line of evidence, in a large-scale finnish study^12^, polygenic predictions were not significant for datasets in which the independence is guaranteed, while in other groups without the guarantee performed significantly higher. Second, we argue that the low portability PRS in trans-ethnic applications may serve as another evidence of overlapping subjects developed for within-ethnic models. Low trans-ethnic portability, yet is not fully understood, has been attributed to different linkage disequilibrium (LD) structures, allele frequencies, and marginal effect size variations according to ancestries. We suspect that the strictly preserved independence of subjects between different ethnicities limits the prediction performance of trans-ethnic models (Fig. 1b). In other words, it is plausibly the level of dependence (i.e., overlapping subjects) between datasets that could explain the gap between within- and trans-ethnic generalization capacities of PRS models.

The aforementioned clues, albeit circumstantial, led us to a systematic investigation for detecting and quantifying the overestimation bias in sPRS due to overlapping subjects. On AD prediction, we first prove that PRS models overfit to overlapping subjects, resulting in overestimated prediction performances on the replication set. Then we extend our experiments using UK Biobank data to derive the scale effect of the inflation and brief guidelines for detecting the bias in sPRS.

## RESULTS

### Overview of the study

Fig. 1 illustrates an overview of our study design. We design an rPRS model that derives SNPs from ADSP, which is then replicated for all subjects from AMP-AD (see Fig. 1b for details and Fig. 2a for results). After removing subjects from AMP-AD that have close kinship with the ADSP study, predictions are made again on AMP-AD, and the two results with and without close subjects are compared. We also compare rPRS and sPRS. To this end, ADSP is divided into 9:1 (discovery:replication) splits for ten-fold cross-validation for rPRS (Fig. 2b), while AMP-AD data are used as another replication set for rPRS (Fig. 2c). An increasing degree of overlapping bias is observed with an expanding number of subjects in the replication set being replaced by samples from the corresponding discovery set (Fig. 2d). We further demonstrate that sPRS also overestimates prediction performances (Fig. 1b, Fig. 2b, and Fig. 2c). We compare the sPRS prediction results against rPRS to indirectly infer the level of overfitting. However, the number of subjects in the IGAP study is larger than that of ADSP, and PRS predictions may not be directly comparable due to the scale effect, where a larger discovery set may result in better generalization capability.

**Fig. 2.**
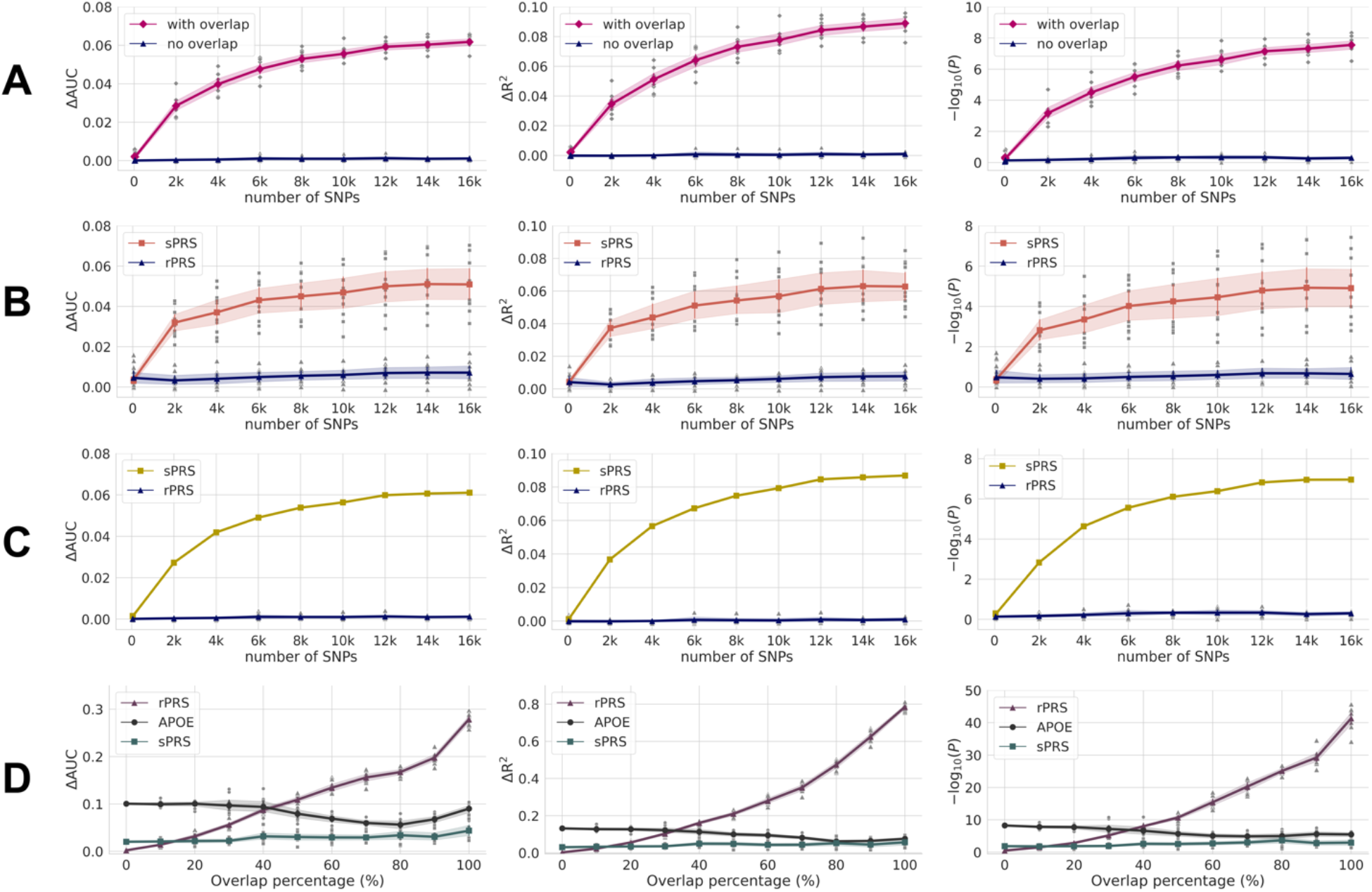
PRS performance comparisons for Alzheimer’s disease. ΔAUC and ΔR^2^ denote the additive gain from introducing PRS term to Model II (refer to Materials and Methods for details). For convenience, we abbreviate the discovery and replication sets as D and R, respectively. (A) AD prediction performances with and without subject overlap (D: ADSP, R: AMP-AD). All metrics of overlapping subjects are overestimated, growing in an increasing number of SNPs. (B) sPRS (D: IGAP, R: ADSP) is compared to rPRS (D: ADSP, R: ADSP). (C) AMP-AD data are another R for rPRS (D: ADSP) and sPRS (D: IGAP). D and R of ADSP data are derived from tenfold cross-validation. In both (B) and (C), sPRS performances are significantly higher than rPRS, and we suspect that some participants of IGAP are identical to a subset of ADSP or AMP-AD. (D) A simulated study is conducted with rPRS (D: ADSP, R: AMP-AD), in which a growing number of subjects in R are replaced by a subset of D (see Results for details). As more subjects overlap between datasets, rPRS performance increases monotonically. Meanwhile, sPRS (D: IGAP, R: the mixture of ADSP and AMP-AD) results remain relatively consistent. Judging by the intersection point of rPRS and sPRS, at least 10% of AMP-AD participants are suspected to overlap with the IGAP study. When only APOE ε4 status is included for developing rPRS in the same manner, performances are relatively stable compared to rPRS, which indicates that the characters of AD are maintained irrespective of the AMP-AD and ADSP combination.

To adjust for the scale effect, a large number of samples in UK Biobank is leveraged in terms of two phenotypes with higher heritabilities than AD^18–20^, namely hypertension and height, which respectively are binary and non-binary variables. With a varying number of subjects in the discovery set, the rate of change per subject in rPRS accuracies is inferred. Finally, we estimate the level of overestimation bias in sPRS for AD prediction (Fig. 1c and Fig. 3).

**Fig. 3.**
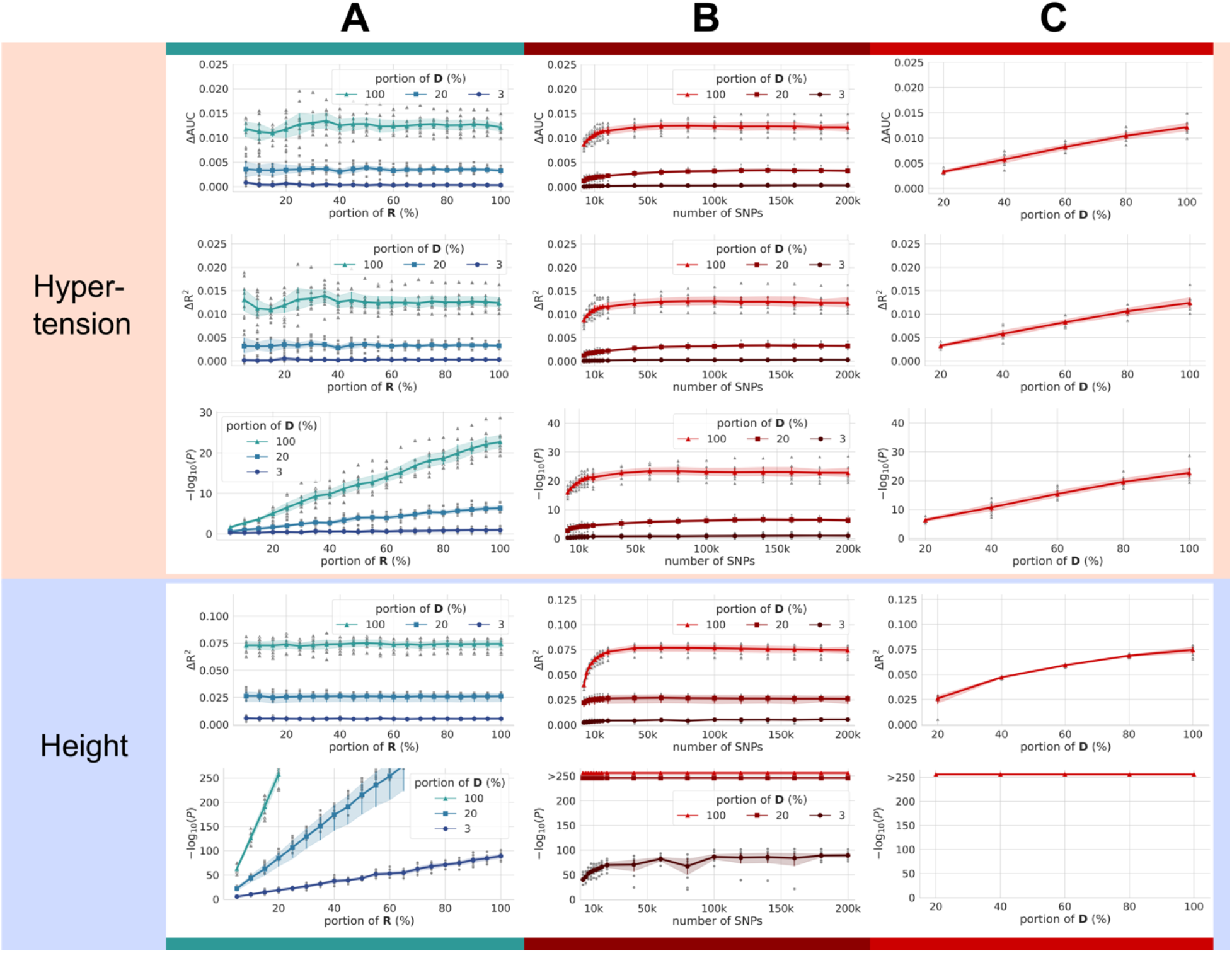
PRS performance comparisons via UK Biobank. In this study, UK Biobank’s primary purpose is to evaluate the scale effect, defined as the marginal gain of performance due to the size of the discovery set. To this end, two variables representative for high heritability, namely hypertension, and height, are analyzed. For experimental purposes, we intentionally design three discovery sets with different sizes, 300k, 60k, and 9k, which approximately correspond to the discovery set sizes of the full UK Biobank dataset, IGAP, and ADSP, respectively. For convenience, we abbreviate the discovery and replication sets as D and R. ΔAUC and ΔR^2^ denote the additive gain from introducing the PRS term to Model II (refer to Materials and Methods for details). (A) A larger D size results in higher prediction performances (ΔAUC and ΔR^2^), demonstrating the scale effect as hypothesized. Yet, in the three sample sizes, a smaller subset of R rarely degrades ΔAUC or ΔR^2^, but it had an impact on the significance level P, perhaps intuitively. As the highest heritability (Fig.1C) foretells, the height variable applied in PRS showed a greater impact on the prediction model than hypertension as indicated by higher ΔR^2^ and −log(P). (B) When the number of SNPs varies with 100% of R used, most metrics show improvements until 50k SNPs are used, after which it plateaus. (C) Although the size of D shows a linear correlation with PRS performances, proving the hypothesized scale effect, the improvements are not dramatic. For instance, ΔR^2^ increases by approximately 0.0000125 and 0.0000083 per 3k of D.

### PRS prediction performance after excluding genetically related individuals

432 identical subjects overlap between ADSP and AMP-AD (Fig. 1b). Using ADSP as the discovery set, rPRS on all subjects in AMP-AD results in ΔAUC of 0.069 (p = 1.51×10^-10^). After removing the overlapping individuals from AMP-AD, the ΔAUC decreases to 0.0017. Notably, ΔAUC loses its statistical significance (p=0.57). ΔR^2^ shows a similar level of deflation, which drops from 0.11 to 0.0041 by removing the identical subjects (Fig. 2a). PRS performances are only slightly affected when close relatives are removed by applying a lower cutoff of PI_HAT, and ΔAUC still shows no statistical significance (Supplementary Table 2).

#### rPRS and sPRS Performances on AD prediction

Fig. 2b, Fig. 2c, and Supplementary Table 3 show the comparison results of rPRS and sPRS. ADSP data are divided into the discovery and replication datasets with 9:1 cross-validation for assessment of rPRS (Fig. 2b). Another test of rPRS is evaluated on non-overlapping data of AMP-AD (n=692) (Fig. 2c). We compute sPRS using summary statistics derived from the first stage of IGAP study (Fig. 2b and Fig. 2c). sPRS on ADSP is evaluated on 10 replication folds (i.e., sets), while, on AMP-AD, the whole data (n=1133) are regarded as a replication set. Unlike rPRS, sPRS is evaluated on all AMP-AD data as overlapping subjects are not identifiable against IGAP.

In rPRS, a respective set (ΔAUC, ΔR^2^) of ADSP and AMP-AD data are (0.0071±0.0052, 0.0077±0.0045) and (0.0013±0.00091, 0.0011±0.0018). P-values are not significant (p>0.05). In other words, when the independence of data is guaranteed, PRS for AD displays unexpectedly low performance. In sPRS, a respective set (ΔAUC, ΔR^2^) of ADSP and AMP-AD are as high as (0.051±0.013, 0.063±0.015) and (0.060, 0.086). The results of sPRS are significantly inflated in comparison to those of rPRS.

### PRS performances are sensitively affected by dependency

Given the suspected inflation in sPRS, we wanted to show how the PRS results are sensitively affected according to the subject overlap, as well as estimate the number of overlapping subjects in sPRS. To this end, we simulate an increasing number of subjects from the discovery set to be added into the replication set. All ADSP data (n=10,923) are used to derive an rPRS model, which subsequently is evaluated on a mixture of AMP-AD (independent replication set, n=692) and ADSP data randomly selected from replication splits of ten-fold cross-validation (fully dependent replication set, n=692), for which the portion of the latter increases from 0 to 100% via an increment of 10%. sPRS is derived from the first stage data of the IGAP study and evaluated in the same way (Fig. 2d). As expected, all ΔAUC, ΔR^2^, and – log(P) monotonically increase in a growing portion of subject overlap. sPRS performances remained relatively unchanged while maintaining the inflated values greater than those in independent rPRS (Fig. 2b and Fig. 2c). To show that the AD characteristics of the replication sets are maintained, performances of *APOE* ε4 are also displayed. Judging by the intersection of two lines representing rPRS and sPRS trends, we infer at least 10% of participants in AMP-AD or ADSP are included in IGAP. However, this holds true only if the number of subjects in discovery sets is equal. Meanwhile, the number of subjects in IGAP is five times larger than that of ADSP. Therefore, to confirm the suspected inflation in sPRS, we must investigate the scale effect of the discovery set.

### Upper bounds of PRS performance derived from UK Biobank data

UK Biobank data are leveraged to infer the upper bounds for the scale effect of sPRS. Hypertension and height are selected as two target variables to investigate the scale effect due to their notably high heritability (Fig. 1c). We posit that AD prediction scores are bounded by both hypertension and height thanks to superior heritability and the scale effect. Subjects from UK Biobank are split by 9:1 following the ten-fold cross-validation scheme.

To investigate the scale effect of AD, rPRS is evaluated on three different sizes of the discovery set, corresponding to 9k, 60k, 300k (i.e., full data) and roughly equal to the size of ADSP, IGAP, and UK Biobank, respectively (Supplementary Table 4). Fig. 3 summarizes and visualizes the results. For hypertension, with a discovery set size of 60k subjects, a set of metrics (ΔAUC, ΔR^2^) is (0.0033±0.00047, 0.0033±0.00051). When the size is 300k, (ΔAUC, ΔR^2^) is (0.012±0.0014, 0.012±0.0017), which is still significantly smaller than the sPRS of ADSP and AMP-AD (Fig. 2b and Fig. 2c) corresponding to (0.051±0.013, 0.063±0.015) and (0.060, 0.086), respectively.

For height, ΔR^2^ for 60k and 300k sizes are 0.028±0.0015 and 0.075±0.0056 (Fig. 3a and Supplementary Table 5). Relatively larger contributions of PRS in height compared to hypertension reflect the greater heritability. In both variables, PRS scores plateau at approximately 50k of SNPs (Fig. 3b). The results of sPRS using IGAP (54k) as a discovery set are highly inflated compared to those of hypertension and height at a similar scale (60k). Scores display growth linear or sublinear to the size of the discovery set (Fig. 3c).

To obtain statistical significance for ΔAUC, a substantially large number of subjects are required for both the discovery and replication sets (Supplementary Fig.) For instance, for a 60k-sized discovery set, more than 10k subjects are needed in the replication set for sufficient power (p<0.01). Lassosum^2^, which uses the LD information, shows two-fold higher performance than PRS (Supplementary Table 6), but the linear pattern of the scale effect remains unchanged.

## DISCUSSION

This study illuminates the overestimation bias in PRS studies. In AD prediction, we prove the presence of latently overlapping subjects for sPRS and demonstrate performance inflation. Developing sPRS without knowledge of the overlapping individuals raises concerns over overestimation and the model’s generalizability. We argue that overestimation should be suspected when the PRS contribution by the referenced PRS methods^21^ exceeds 1–2% of ΔR^2^ for binary traits^17^ and 7–8% of ΔR^2^ for non-binary traits, which are the respective upper limits drawn by hypertension and height from UK Biobank.

As evidenced by the three separate AD studies sharing identical subjects, subject overlap may be prevalent in other cohort studies and large-scale meta-analyses. In our study, ADSP and AMP-AD data include 432 identical subjects, which corresponds to 38.12% of AMP-AD data. Also, ADSP and ADNI have 21 overlapping subjects (Fig. 1a). As genetic studies are conducted across multiple centers, the odds of having duplicated subjects are high among different initiatives. Therefore, planning of PRS studies requires considerable attention to exclude identical persons.

Several existing studies support the relationship between the independence of datasets and the possibility of the overestimation bias. Concretely, we observe the trend in which PRS is performing significantly lower when the independence of data is explicitly controlled. For instance, when overlapping subjects are removed, PRS contributed less than 2% in accuracy even with large discovery sets from a national biobank ^12,17^. In a large-scale finnish study for coronary artery disease (CAD), arterial fibrillation (AF), type 2 diabetes mellitus (T2DM), breast cancer (BrC) and prostate cancer (PrC), the independencies in CAD and AF were clarified^12^. The PRS contributions of two diseases were statistically insignificant (0.3% and 0.9%, respectively). In contrast, the PRSs of T2DM, BrC, and PrC used the summary statistics derived from large meta-analyses which included finnish population, the contributions of PRS were significantly high (2%, 3.9%, and 2.9%, respectively).

We propose two potential signs when the overestimation bias should be suspected. The first is when a PRS model achieves surprisingly high performance without sufficient participants. The number of subjects and SNP heritability are important factors in PRS performance^18^. For example, hundreds of thousands of subjects in the discovery set would be required for PRS to be used in disease prediction^18^. Computing the PRS using UK Biobank for traits with prominent heritability sets the upper bounds for other traits with lower heritability. Thus, we reveal that a replication set of 10k is required in the 60k discovery set for statistically significant for AUC increment by PRS. In the studies registered in the PGS catalog, the median number of subjects in the replication sets is 6995 and 24,573 in the discovery sets^22^. In addition, a systematic review of AD PRS studies reveals that the replication set size ranges from 59 to 116,666^23^. About one third of them, the sample size is less than a thousand. Therefore, an abundance of prior sPRS studies^23–25^ may not suffice as the number of samples required for statistically significant results.

Second, for particular phenotypes, high variability in performances across different nations or races using the same discovery set may be another sign of overestimation. PRS performances plummet if the discovery and replication sets are from different countries or ethnicities^12–16^. However, a line of evidence suggests that trans-ethnic portability remains in many traits^26–28^, even if the inherent differences in LD structures across races prevent causal variants from being correctly reflected in PRS^29^. For instance, Martin et al. reported equivalent performances within the confidence interval for intra- and trans-ethnic replication sets in four out of five binary traits and 11 out of 17 non-binary traits of BioBank Japan^26^. Also, sPRS developed with Europeans showed a 50% discounted performance for East Asians and 25% for Africans^26^. Therefore, the gap (or the variability) between intra-ethnic and trans-ethnic evaluations of PRS is attributable to overestimation bias, rather than a low trans-ethnic portability. For example, if the performance for a trans-ethnic application is preserved at less than 25% for intra-ethnic evaluation, overlapping bias might be suspected. In other words, sPRS suffers in trans-nation or trans-ethnic studies, since the subject-level independence strictly holds.

PRS also could be developed using GWA (p<5×10^-8^) SNPs in core genes curated from multiple GWA studies^30–32^, which we argue are not free from the overestimation bias. Using GWA SNPs is justified because it substitutes the P-value thresholding step required in conventional PRS studies for selecting SNPs. Also, GWA SNPs tend to show highly significant P-values, and therefore are regarded as reliable. Moreover, GWA SNPs information can be conveniently accessible via reviewing prior works, even if the authors do not release summary statistics. However, GWA SNPs are frequently found in the uninterpretable non-coding regions^33^, and PRS performances increase with a higher number of SNPs then plateau (Fig. 2 and Fig. 3)^34^. Therefore, PRS models from GWA SNPs may overfit to the discovery set. Hence, it follows that the odds of overestimation bias are high when the GWA SNPs are selected from multiple studies.

For the non-binary trait height, PRS has a greater contribution than binary trait hypertension. Similar trends have been observed in a prior study^26^. Heritability for height and hypertension was reported as 49.7% and 14.7%, respectively^20^. Although a superior heritability of height partially explains the performance gap, characteristics of each phenotype may also play a role. For instance, while measuring height is straightforward, a diagnosis of hypertension can depend on age. Thus, a subset of the control group can later be diagnosed with hypertension.

One limitation in our study is the indirect derivation of the scale effect to prove the overestimation bias of sPRS in AD prediction. We justify the choice of our methods based on three reasons. First, hypertension and height have higher heritabilities than AD^19,20,30^. Second, the number of SNPs used for UK Biobank is an order of magnitude larger than those used for AD data analysis. Finally, the number of available subjects in UK Biobank is five-fold larger than IGAP. Therefore, we argue that PRS performances of hypertension from UK Biobank are sufficient upper bounds not only for AD prediction but also for most binary complex genetic traits/diseases.

As the risk of overestimation bias is evaluated in sPRS studies, care must be taken to prevent overlap, especially within the same ethnicity. While applications of genetic studies continue to gain momentum and many countries create large-scale biobanks^35–38^, PRS developed from large meta-analyses that curate and merge data from several countries should be screened in advance to filter out overlapping subjects. Researchers often release summary statistics to help further research^22^. Direct comparisons between a replication set and large-scale data are difficult when using summary statistics. As such, we showcase both direct and indirect methods to probe overestimation bias within the same ethnicity—either of which, we argue, must be mandatory to improve PRS reliability.

## METHODS

### Participants

In this work, AD genetic studies–International Genomics of Alzheimer’s Project (IGAP), Alzheimer’s Disease Sequencing Project (ADSP), and AMP-AD–are used to demonstrate overestimation bias in PRS^39–45^. Non-Hispanic white individuals are used, and their cross-study genetic relatedness is revealed by principal component (PC) and identity-by-state analyses. After quality control, our final analyses include 10,293 participants from ADSP and 1,133 from - AD (Supplementary Table 1). In the UK Biobank database^46^, a total of 342,318 white-British participants had hypertension and height records. Refer to the Supplementary Material for additional information about the study datasets, sequencing methods, and quality control processes.

### Statistical analyses

We perform logistic regressions for binary traits and linear regressions for a continuous phenotype, using PLINK (v1.9)^47^. Three different regression models are constructed. First, a simple regression model is used with PRS as the only covariate. Second, Model II denotes a multivariable regression without PRS, consisting of additional covariates highly related to the phenotypes. In Model III, we introduce PRS as an additional covariate to Model II. In both models, we control for 20 leading PCs

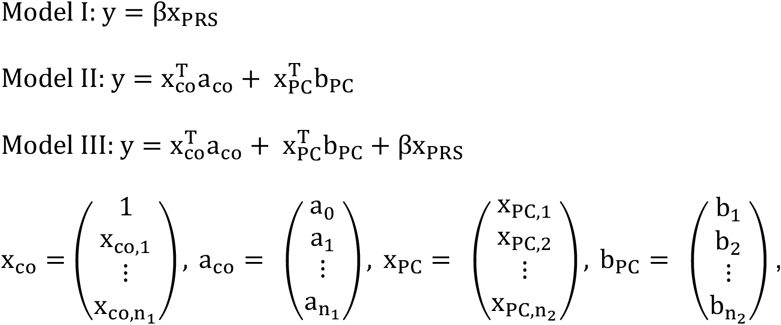

Where x_co_, 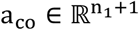 are vectors of general covariates (e.g., age and sex) and corresponding coefficients, respectively, while x_PC_, 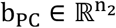 are vectors of PCs and PC coefficients, respectively. Here, x_PRS_ denotes the PRS term. Throughout the manuscript, we focus on measuring the additive gain of PRS in Model III, on top of Model II.

For AD datasets, common covariates include sex, APOE ε4 status, and the sequencing centers. PCs are computed using the principal component analysis function of PLINK (v1.9)^47^. For UK Biobank, age, sex, and array types (UK Biobank Axiom array or UK BiLEVE Axiom array) are considered as covariates. Here, we download 40 PCs pre-calculated with fastPCA^48^. For hypertension of UK Biobank, body mass index additionally is included in the covariates. For binary traits, the areas under receiver operating characteristic (AUC) and Nagelkerke’s pseudo-R^2^ are used to assess model performances, which are calculated using “pROC” and “fsmb” packages of R (v4.0.3), respectively^49,50^. Performance improvements from PRS are determined by subtracting AUC and R^2^ of Model II from those of Model III, which we respectively label as ΔAUC and ΔR^2^. The statistical significance of ΔAUC is examined using DeLong’s test^51^. For height, a non-binary trait, we compute the adjusted-R^2^ via “lm” in the R program. PRS contributions are determined by comparing Model II and Model III with the extra sum of squares test.

### Cross-validation

For ten-fold cross-validation tests, we balance the number of samples between the discovery and replication splits based on each covariate in the statistical analyses using “StratifiedKFold” function from Python’s (v3.8) “scikit-learn” (v0.24.1) package^52^. When testing a part of the cross-validated datasets, samples are balanced over covariates using the R (v4.0.3) “sampling” (v2.9) package^53^.

### Computation of PRS

For computing PRS, we select common (MAF≥1%) SNPs and use summary statistics from discovery sets, followed by measuring PRS in replication datasets. After selecting SNPs with p<0.5 in the association tests using the discovery dataset, we perform clumping with the window of ±1Mbp and r^2^<0.1. Clumping is performed using PLINK (v1.9)^47^. For AD genetic studies, we exclude any SNPs within 1Mbp of the *APOE* (apolipoprotein E) region. When analyzing the effect of the number of SNPs on the results, the SNPs are selected in the order from the lowest P-value. We construct PRS with PRSice (v2.3) and Lassosum (v0.4.5)^2,21^.

## COMETING INTERESTS

The authors declare that they have no competing interests.

## DATA AVAILABILITY

The dataset(s) supporting the conclusions of this article are available in webpages of UK Biobank (https://www.ukbiobank.ac.uk/), ADSP (accession phs000572.v1.p1; https://www.ncbi.nlm.nih.gov/projects/gap/cgi-bin/study.cgi?study_id=phs000572.v1.p1), Mayo RNAseq study (accession syn5550404; http://dx.doi.org/10.1038/sdata.2016.89), Mount Sinai Brain Bank (MSBB) study (accession syn3159438; http://dx.doi.org/10.1038/sdata.2018.185), and Religious Orders Study and Memory and Aging Project (ROSMAP) Study (accession syn3159438; http://dx.doi.org/10.1038/mp.2017.20).

## AUTHOR CONTRIBUTIONS

D.K.P., M.C., S.Y., and J.H.K. conceived and designed the study. D.K.P. and J.H.K. performed statistical analysis. D.K.P., M.C., and J.H.K. analyzed the genetic data. All authors discussed the results and implications and commented on the manuscript at all stages. M.C., S.K., Y.Y.J., R.K.L., H.S.K., J.C., and S.Y. gave technical support and conceptual advice. D.K.P., S.Y., and J.H.K. wrote the paper. All co-authors contributed to the final manuscript.

