## Supplementary methods, tables, and figure for "Overestimated Polygenic Prediction due to Overlapping Subjects in Genetic Datasets"

Contents

1. Supplementary Methods
   - 1.1 UK Biobank data
   - 1.2 International Genomics of Alzheimer's Project (IGAP)
   - 1.3 ADSP and AMP-AD data
   - 1.4 Genetic relatedness
   - 1.5 Data processing and genotype QC of the UK Biobank
   - 1.6 Data processing and genotype QC of the ADSP study
   - 1.7 Data processing and genotype QC of the AMP-AD study
   - 1.8 Imputation and selection of SNPs

2. Supplementary Tables and Figures
   - Supplementary Table 1. Demographic characteristics
   - Supplementary Table 2. PRS performance after excluding genetically close individuals from the discovery set
   - Supplementary Table 3. Results of rPRSs and sPRS
   - Supplementary Table 4. PRS performances of hypertension in UK Biobank
   - Supplementary Table 5. PRS performances of height in UK Biobank
   - Supplementary Table 6. Comparison between two PRSs methods
   - Supplementary Figure. Least number of replication set subjects required to gain statistical significance (p<0.01) for hypertension using UK Biobank

3. References

1. Supplementary Methods

1.1 UK Biobank data

The UK Biobank recruited approximately 500,000 individuals in 22 centers of the United Kingdom from 2006 to 2010[1]. The recruited people ranged in age from 38- to 73-years-old. Extensive phenome information for laboratory findings, physical measurements, health records, and questionnaires about health and lifestyles were collected. We use the self-reported status of hypertension and the standing height at the baseline enrollment. Detailed descriptions are in the previous landmark papers of UK Biobank[1,2]. Data are available for download from www.ukbiobank.ac.uk.

1.2 International Genomics of Alzheimer's Project (IGAP)

Stage 1 of IGAP used the four previously published Genome-Wide Association Study (GWAS) datasets: EADI (European Alzheimer's Disease Initiative) [3,4], ADGC (Alzheimer's Disease Genetics Consortium) [5], CHARGE (The Cohorts for Heart and Aging Research in Genomic Epidemiology Consortium) [6], and GERAD (Genetic and Environmental Risk in AD Consortium) [7]. Stage 1 includes 17,008 Alzheimer's disease (AD) cases and 37,154 controls. Stage 2 includes 8,572 AD cases and 11,312 controls. All participants are individuals of European ancestry. Finally, a meta-analysis is performed, combining results from Stages 1 and 2[8]. In this study, the summary statistics of Stage 1 were downloaded from http://web.pasteur-lille.fr/en/recherche/u744/igap/igap_download.php.

1.3 Alzheimer's Disease Sequencing Project (ADSP) and AMP-AD data

ADSP is a whole-exome sequencing study that consists of 5,844 AD cases and 4,767 controls. AMP-AD is a whole-genome sequencing study with 955 AD cases and 821 controls. The participants of ADSP are derived from 24 cohorts of AD Genetics Consortium and cohorts for Heart & Aging Research in Genomic Epidemiology. All cases meet the NINCDS-ADRDA (National Institute of Neurological and Communicative Diseases and Stroke/Alzheimer's Disease and Related Disorders Association) criteria for possible, probable, or definite AD with documented age at onset or age at death (for pathologically verified cases), as well as *APOE* (Apolipoprotein E) genotyping. All controls are at least 60-years-old and free of dementia by direct or documented cognitive assessment. Additional descriptions of the sample selection criteria for the ADSP samples can be found elsewhere[9]. The AMP-AD study consists of three genetics studies: The Mayo Clinic AD Genetics Studies[10], Mount Sinai Brain Bank (MSBB) study[11], and Religious Orders Study and Memory and Aging Project (ROSMAP) Study[12].

1.4 Genetic relatedness

We exclude participants with closer kinship within the third degree using the kinship matrix precalculated and deposited in the UK Biobank webpage. The kinship matrix has been calculated using KING[13]. For AD genetic studies, we obtained PI_HAT values using the genome function of PLINK (v1.9) after merging the common variants (MAF≥1%) of ADSP and AMP-AD data[14]. Before merging, the participants with close relatedness (PI_HAT > 0.1 in ADSP and PI_HAT > 0.2 in AMP-AD) within the studies are excluded. Subjects from AMP-AD are regarded as genetically identical to those in ADSP when PI_HAT=1.0 (n=432) and closely related when PI_HAT>0.2 (n=441).

1.5 Data processing and genotype QC of the UK Biobank

Using the UK BiLEVE array and UK Biobank axiom array, 488,377 participants' samples are successfully genotyped. We involve only the individuals with white-British ancestry in this study. We exclude the third-degree related individuals (kinship coefficients < 0.0442). Also, individuals are excluded when there is a sex mismatch or sex chromosome aneuploidy, or there are outliers in heterozygosity and missing rates. Finally, a total of 342,318 individuals who have hypertension and height phenotypes are used in the study.

1.6 Data processing and genotype QC of the ADSP study

Sequencing for the ADSP project was performed in three centers: the Broad Institute, Human Genome Sequencing Center at the Baylor College of Medicine, and McDonnell Genome Institute at Washington University in St. Louis. The quality control (QC) step of the sequenced whole-exome sequencing (WES) data followed the GATK (genomic analysis toolkit) best practices. The inclusion criteria for variants are mapping score ≥0.8, high (≥10) mean read depth, in-range VR/DP (variant read/total read depth ratio ≥0.25 and ≤0.75), and allele balance (AB) scores ranging from 0.3 to 0.7. The exclusion criteria for variants are high (≥20%) missingness, very high (≥500) read depth, excess heterozygosity (for MAF<0.2, |z|>1.22; for MAF≥0.2,

|z|>5SD), monomorphic variants, variants with controls' deviation of Hardy-Weinberg Equilibrium ($p<5.0\times10^{-5}$, only in MAF>0.1%), and variants with missingness test $p<1.0\times10^{-8}$ between cases and controls. The exclusion criteria for individuals are excess (>6SD) private variants, low (≤80%) call rate, high (>6SD) Ti/Tv ratio, high (>6SD) heterozygosity-to-homozygosity ratio, and high (>500) mean read depth. Finally, 10,907 subjects with 1,508,430 variants in a case-control study remained. We exclude outliers using the leading two principal components (PCs). Among them, we analyzed the data of 10,294 European participants who have phenotypic information.

1.7 Data processing and genotype QC of the AMP-AD study

After downloading whole-genome sequencing data (WGS) from https://adknowledgeportal.synapse.org/, we select variants with high tranche sensitivity (>99.9%) and for their quality (≥30). Additionally, the genotypes with high-quality depth (QD≥5 for indels and QD≥2 for single nucleotide polymorphisms [SNPs]), in-range AB (>0.3 and <0.7), high (≥5) mean depth, and low (≤5%) missingness are selected. We exclude variants with maximal read depth >5SD and those in the low complexity regions. We discard variants with high significance ($p<1.0\times10^{-8}$) in the missingness tests (between cases and control; different studies) and deviation of Hardy-Weinberg Equilibrium ($p<1.0\times10^{-6}$) in controls. The individuals with high (>10%) missing genotypes are excluded. We check sex with PLINK (v1.9) [14].

1.8 Imputation and selection of SNPs

Version 3 imputation genotype data are downloaded from www.ukbiobank.ac.uk. Before imputation, the genotyped data are phased using a modified version of the SHAPEIT2 algorithm[15]. The multiallelic or rare (MAF ≤1%) SNPs are removed before phasing. Genotype imputation is performed using IMPUTE2 algorithms[16]. The UK10K haplotype and HRC reference panels are combined for the imputation reference set[17]. The imputation INFO cutoff scores differ according to MAFs (Info>0.3 for MAF >3%; Info>0.6 for MAF 1-3%; Info>0.8 for MAF 0.5-1%; Info>0.9 for MAF 0.1-0.5%) [1]. We filter out SNPs with MAF < 1% and deviation of Hardy-Weinberg Equilibrium ($p<1.0\times10^{-8}$).

To increase the common variants between ADSP and AMP-AD data, we use imputed data. After selecting variants with MAF ≥ 1% and call rate > 95% in addition to genotype QC, we perform imputation using the Michigan imputation server (https://imputationserver.sph.umich.edu/) with multi-population reference panels of Phase III 1000 genomes and default parameters. After imputation, we discard variants with MAF < 1% and $R^2 < 0.3$.

Supplementary Table 1. Demographic characteristics

(A) ADSP and AMP-AD

| | ADSP | | AMP-AD | |
|---|---|---|---|---|
| | Cases n = 5,687 | Controls n = 4,606 | Cases n = 696 | Controls n = 437 |
| Age: years | 75.51±8.83[a] | 87.21±4.78[a] | 85.07±6.08 | 82.34±7.20 |
| Women No. (%) | 3,267 (57) | 2,698 (59) | 466 (67) | 267 (61) |
| *APOE* ε4 No. (%) | 2,405 (42) | 647 (14) | 254 (36) | 68 (15) |

(B) UK biobank

| | Hypertension | |
|---|---|---|
| | Cases n = 82,719 | Controls n = 259,599 |
| Age: years | 59.70±6.91[a] | 55.97±8.10[a] |
| Women No. (%) | 37,844 (46) | 146,128 (56) |
| BiLEVE Axiom[b] No. (%) | 9,860 (12) | 27,466 (11) |
| Height (m) | 1.69±0.09[a] | 1.69±0.09[a] |

In cases of ADSP, age means the age at onset, while in controls of ADSP, it is age at the last assessment. The age and height of UK Biobank are those at the baseline enrollment

[a] means ± standard deviations
[b] There are two DNA chip platforms: Affymetrix UK BiLEVE Axiom or Affymetrix UK Biobank Axiom array

Supplementary Table 2. PRS performance after excluding genetically cloase individuals from the discovery set

| | | AUC | | | | $R^2$ | | | | |
|---|---|---|---|---|---|---|---|---|---|---|
| PI_HAT[a] | number[b] | Model I | Model II | Model III[c] | ΔAUC[c] | Model I | Model II | Model III | $\Delta R^2$ | P-value[d] |
| | 1133 | 64.78 | 74.03 | 80.91 | 0.069 | 0.069 | 0.14 | 0.26 | 0.11 | $1.51\times10^{-10}$ |
| 1.0 | 701 | 51.27 | 80.53 | 80.70 | 0.0017 | –0.0014 | 0.23 | 0.24 | 0.0041 | 0.57 |
| 0.9 | 697 | 51.35 | 80.68 | 80.79 | 0.0012 | –0.0014 | 0.24 | 0.24 | 0.0033 | 0.67 |
| 0.8 | 697 | 51.35 | 80.68 | 80.79 | 0.0012 | –0.0014 | 0.24 | 0.24 | 0.0033 | 0.67 |
| 0.7 | 697 | 51.35 | 80.68 | 80.79 | 0.0012 | –0.0014 | 0.24 | 0.24 | 0.0033 | 0.67 |
| 0.6 | 697 | 51.35 | 80.68 | 80.79 | 0.0012 | –0.0014 | 0.24 | 0.24 | 0.0033 | 0.67 |
| 0.5 | 694 | 51.17 | 80.60 | 80.73 | 0.0013 | –0.0014 | 0.24 | 0.24 | 0.0036 | 0.65 |
| 0.4 | 692 | 51.20 | 80.67 | 80.80 | 0.0013 | –0.0014 | 0.24 | 0.24 | 0.0032 | 0.64 |
| 0.3 | 692 | 51.20 | 80.67 | 80.80 | 0.0013 | –0.0014 | 0.24 | 0.24 | 0.0032 | 0.64 |
| 0.2 | 692 | 51.20 | 80.67 | 80.80 | 0.0013 | –0.0014 | 0.24 | 0.24 | 0.0032 | 0.64 |
| 0.1 | 692 | 51.20 | 80.67 | 80.80 | 0.0013 | –0.0014 | 0.24 | 0.24 | 0.0032 | 0.64 |

[a] Calculated after merging ADSP and AMP-AD
[b] The number of participants of AMP-AD after excluding individuals ≥ PI_HAT
[c] Refer to “Material and Methods” for models
[d] P-value for differences of AUC of Model III and Model II
Abbreviation: AUC, area under curve

Supplementary Table 3. rPRS and sPRS results on AD.

| | Dataset | | AUC | | | | $R^2$ | | | | |
|---|---|---|---|---|---|---|---|---|---|---|---|
| | Discovery | Replication | Model I | Model II[b] | Model III | ΔAUC[c] | Model I | Model II | Model III | $\Delta R^{2c}$ | –log(p)[d] |
| rPRS | ADSP[a] (n=9k) | ADSP[a] (n=1k) | 0.56±0.013 | 0.70±0.017 | 0.70±0.018 | 0.0071±0.0052 | 0.0098±0.0049 | 0.10±0.019 | 0.11±0.021 | 0.0077±0.0045 | 0.65±0.47 |
| | ADSP[a] (n=9k) | AMP-AD[e] (n=0.7k) | 0.51±0.019 | 0.79 | 0.79±0.00091 | 0.0013±0.00091 | –0.0011±0.00032 | 0.20 | 0.20±0.0018 | 0.0011±0.0018 | 0.35±0.14 |
| sPRS | IGAP (n=54k) | ADSP[a] (n=1k) | 0.66±0.012 | 0.70±0.017 | 0.75±0.014 | 0.051±0.013 | 0.076±0.012 | 0.10±0.019 | 0.16±0.022 | 0.063±0.015 | 4.94±1.61 |
| | IGAP (n=54k) | AMP-AD[e] (n=0.7k) | 0.65 | 0.79 | 0.81 | 0.020 | 0.061 | 0.20 | 0.23 | 0.031 | 1.85 |
| | IGAP (n=54k) | AMP-AD[f] (n=1.1k) | 0.69 | 0.72 | 0.78 | 0.060 | 0.11 | 0.12 | 0.20 | 0.086 | 6.87 |

These results are graphically outlined in Fig. 2b and Fig. 2c of the main manuscript
After clumping within 1Mbp, all SNPs with $p < 0.5$ are used, and the number of SNPs is ~17K

[a] Discovery and replication datasets of ADSP are independent and split based on ten-fold cross-validation
[b] Because results from Model II depend only on the replication dataset and not the discovery dataset, they are identical for a particular replication dataset
[c] The additive difference in the performance of Model III against Model II
[d] p values are computed by comparing the AUC of both Model II and Model III via Delong's methods[18].
[e] AMP-AD data without a close kinship with ADSP data
[f] All AMP-AD data
Abbreviations: AUC, the area under the curve; PC, principal component; PRS, polygenic risk score

Supplementary Table 4. PRS performance comparisons for hypertension in UK Biobank.

(A) By the number of subjects in the discovery set

| No of Discovery dataset | AUC | | | | $R^2$ | | | | |
|---|---|---|---|---|---|---|---|---|---|
| | Model I | Model II | Model III | ΔAUC | Model I | Model III | Model II | $\Delta R^2$ | –log(p) |
| 9k | 0.51±0.0060 | 0.70±0.0044 | 0.70±0.0043 | 0.00034±0.00023 | 0.00025±0.00017 | 0.094±0.0041 | 0.095±0.0040 | 0.00029±0.00018 | 0.96±0.64 |
| 60k | 0.54±0.0039 | 0.70±0.0044 | 0.71±0.0042 | 0.0033±0.00047 | 0.0030±0.00060 | 0.094±0.0041 | 0.098±0.0041 | 0.0033±0.00051 | 6.36±0.89 |
| 300k | 0.57±0.0037 | 0.70±0.0044 | 0.72±0.0041 | 0.012±0.0014 | 0.011±0.0011 | 0.094±0.0041 | 0.11±0.0041 | 0.012±0.0017 | 22.69±2.65 |

The number of subjects in the replication set is fixed at 34k

(B) By the number of subjects in the replication set

| No of Discovery dataset | AUC | | | | $R^2$ | | | | |
|---|---|---|---|---|---|---|---|---|---|
| | Model I | Model II | Model III | ΔAUC | Model I | Model III | Model II | $\Delta R^2$ | –log(p) |
| 1.7k (5%) | 0.53±0.0130 | 0.72±0.014 | 0.73±0.012 | 0.0036±0.0027 | 0.0030±0.0022 | 0.091±0.019 | 0.094±0.018 | 0.0032±0.0028 | 0.60±0.37 |
| 17k (50%) | 0.57±0.0054 | 0.70±0.0050 | 0.70±0.0043 | 0.0039±0.00080 | 0.0033±0.00080 | 0.089±0.0045 | 0.092±0.0044 | 0.0036±0.00080 | 3.98±0.78 |
| 31k (90%)[a] | 0.57±0.0047 | 0.70±0.0041 | 0.70±0.0041 | 0.0035±0.00066 | 0.0031±0.00076 | 0.090±0.0038 | 0.093±0.0038 | 0.0034±0.00063 | 5.98±1.09 |

The number of subjects in the discovery set is fixed at 60K, roughly equivalent to the number of the first stage of IGAP
These results are graphically outlined in Fig. 3 of the main manuscript
After clumping within 1Mbp, all SNPs with $p < 0.5$ are used, and the number of SNPs is ~218k

[a] The results of the 100% replication set are in the second row of Supplementary Table 2A.

Supplementary Table 5. PRS performance comparisons for height in UK Biobank
(A) By the number of subjects in the discovery set

| No of Discovery dataset* | $R^2$ | | | | |
|---|---|---|---|---|---|
| | Model I | Model II | Model III | $\Delta R^2$ | –log(p) |
| 9k | 0.0058±0.00036 | 0.54±0.0019 | 0.54±0.0022 | 0.0054±0.00047 | 88.84±7.76 |
| 60k | 0.030±0.0021 | 0.54±0.0019 | 0.56±0.0022 | 0.028±0.0015 | ∞ |
| 300k | 0.077±0.0063 | 0.54±0.0019 | 0.61±0.0061 | 0.075±0.0056 | ∞ |

The number of subjects in the replication set is fixed at 34k

(B) By the number of subjects in the replication set

| No of Replication dataset | $R^2$ | | | | |
|---|---|---|---|---|---|
| | Model I | Model III | Model II | $\Delta R^2$ | –log(p) |
| 1.7k (5%) | 0.032±0.0059 | 0.54±0.013 | 0.57±0.018 | 0.029±0.0037 | 23.99±2.67 |
| 17k (50%) | 0.030±0.0021 | 0.54±0.0043 | 0.56±0.0045 | 0.028±0.0014 | 234.61±11.81 |
| 31k (90%) | 0.030±0.0022 | 0.54±0.0026 | 0.56±0.0024 | 0.028±0.0015 | ∞ |

The number of subjects in the discovery set is fixed at 60k

These results are graphically outlined in Fig. 3 of the main manuscript
After clumping within 1Mbp, all SNPs with p < 0.5 are used, and the number of SNPs is ~221K

Supplementary Table 6. Performance comparisons between PRS [19] and Lassosum [20]

(A) Hypertension

| No of Discovery dataset | ΔAUC | | $\Delta R^2$ | | –log(p) |
|---|---|---|---|---|---|
| | PRS | Lassosum | PRS | Lassosum | |
| 9k | 0.00031±0.00022 | 0.00065±0.00046 | 0.00028±0.00018 | 0.00060±0.00042 | 1.10±0.87 |
| 60k | 0.0033±0.00047 | 0.0063±0.0018 | 0.0033±0.00051 | 0.0064±0.0017 | 6.34±2.84 |
| 300k | 0.012±0.0014 | 0.026±0.0016 | 0.012±0.0018 | 0.028±0.0018 | 23.61±3.18 |

(B) Height

| No of Discovery dataset | $\Delta R^{2}$[a] | |
|---|---|---|
| | PRS | Lassosum |
| 9k | 0.0054±0.00047 | 0.011±0.00071 |
| 60k | 0.028±0.0015 | 0.065±0.0055 |
| 300k | 0.075±0.0056 | 0.12±0.0090 |

[a] Extra sum of squares tests used for comparing Model II and Model III could not calculate the P-values for comparison of models between PRS and Lassosum

Supplementary Fig. Number of replication set subjects required to gain statistical significance ($p<0.01$) for hypertension using UK Biobank.
P-values are obtained by comparing AUC of Model II and Model III. The five lines denote different sizes of the discovery set (D), where 100% corresponds to 300k. The x-axis shows the size of the replication set (R) where 100% is 34k. The small plot in the left upper quadrant shows a higher magnification of the red box area, in which the significance threshold is drawn using a dotted red line

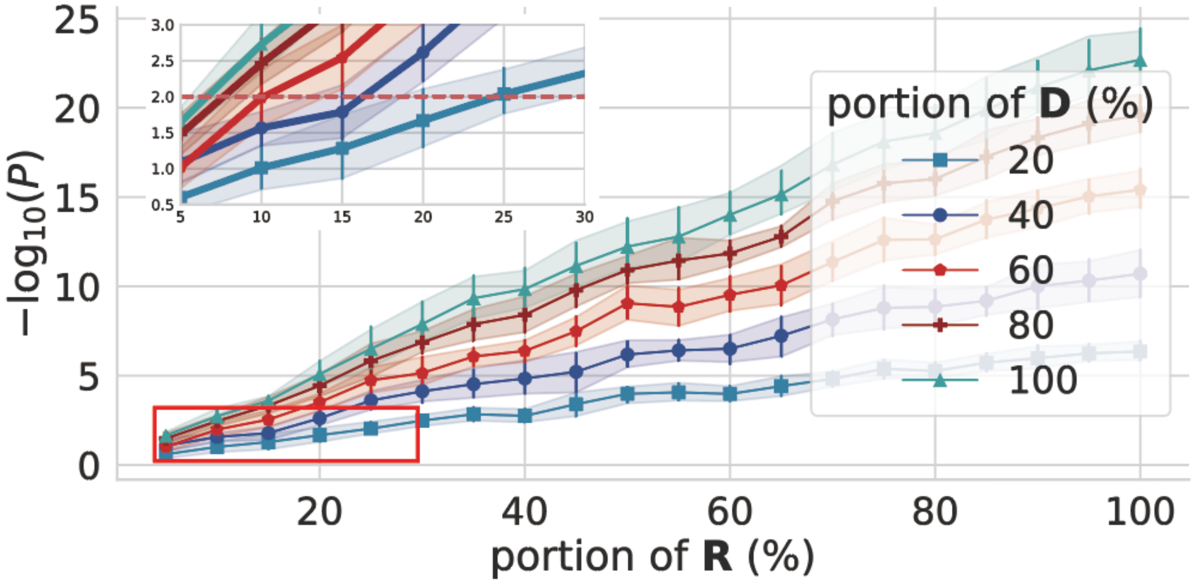
